# Fine-scale quantification of GC-biased gene conversion intensity in mammals

**DOI:** 10.1101/2021.05.05.442789

**Authors:** Nicolas Galtier

## Abstract

GC-biased gene conversion (gBGC) is a molecular evolutionary force that favours GC over AT alleles irrespective of their fitness effect. Quantifying the variation in time and across genomes of its intensity is key to properly interpret patterns of molecular evolution. In particular, the existing literature is unclear regarding the relationship between gBGC strength and species effective population size, *N*_*e*_. Here we analysed the nucleotide substitution pattern in coding sequences of closely related species of mammals, thus accessing a high resolution map of the intensity of gBGC. Our maximum likelihood approach shows that gBGC is pervasive, highly variable among species and genes, and of strength positively correlated with *N*_*e*_ in mammals. We estimate that gBGC explains up to 60% of the total amount of synonymous AT*→*GC substitutions. We show that the fine-scale analysis of gBGC-induced nucleotide substitutions has the potential to inform on various aspects of molecular evolution, such as the distribution of fitness effects of mutations and the dynamics of recombination hotspots.

## Introduction

GC-biased gene conversion (gBGC) is a recombination-associated transmission bias by which G and C alleles are favoured over A and T alleles. This evolutionary force was discovered in the 2000’s from the analyses of early population genomic data sets (Eyre-Walker, 1999; Galtier, Piganeau, et al., 2001; Spencer et al., 2006; Webster and NG Smith, 2004), and experimentally confirmed later on (Mancera et al., 2008; Pratto et al., 2014; Williams et al., 2015). gBGC manifests itself as a GC-bias that affects both non-functional and functional sequences and is correlated with the local recombination rate (Galtier and Duret, 2007). gBGC has a strong impact on patterns of variation genome wide in mammals (Clément and Arndt, 2011; Duret and Arndt, 2008; Pracana et al., 2020; Romiguier, Ranwez, et al., 2010) and many other taxa (Clément, Sarah, et al., 2017; Figuet, Ballenghien, Romiguier, et al., 2014; Galtier, Roux, et al., 2018; Lassalle et al., 2015; Long et al., 2018; Mugal, Arndt, et al., 2013; Nabholz et al., 2011; Pessia et al., 2012; Wallberg et al., 2015). gBGC can mimic the effect of natural selection and confound its detection by generating patterns of clustered AT*→*GC substitutions, distorted site frequency spectra and altered non-synonymous/synonymous ratios (Bolívar et al., 2018; Corcoran et al., 2017; Dreszer et al., 2007; Galtier and Duret, 2007; Lartillot, 2012; Ratnakumar et al., 2010; Rousselle et al., 2019). Importantly, because it favours G and C alleles irrespective of their fitness effect, gBGC tends to counteract natural selection and increase the deleterious mutation load (Berglund et al., 2009; Galtier, Duret, et al., 2009; Lachance and Tishkoff, 2014; Necşulea et al., 2011).

The abundant body of literature reviewed above demonstrates a significant effect of gBGC in a large number of genomes. Only a few studies, however, have attempted to quantify its strength - a harder task. gBGC results from a DNA repair bias involving paired chromosomes at meiosis, and operating in the immediate neighborhood of DNA double strand breaks. The genome average transmission bias, *b*, is therefore expected to be proportional to the recombination rate, gene conversion tract length, and repair bias. The effect of gBGC on genome evolution is also expected to be dependent on the intensity of drift: being a directional force, gBGC is only effective if stronger than the stochastic component of allele frequency evolution. The intensity of drift is inversely related to the effective population size *N*_*e*_, so that the strength of gBGC is usually measured by the *B* = 4*N*_*e*_*b* parameter. Glémin et al. (2015) used genome-wide resequencing data to estimate *B* at the megabase scale throughout the human genome. Fitting various population genetic models to polarised GC vs. AT site frequency spectra, Glémin et al. (2015) estimated the genome average *B* to be in the weak selection range, around 0.4, with *B* reaching a value above 5 in 1%-2% of the genome. This variance among genomic regions in gBGC strength is interpreted as reflecting the existence of recombination hotpots in humans (Capra et al., 2013; Duret and Arndt, 2008; Spencer et al., 2006).

Similar analyses have been performed in a number of non-human taxa. In the fruit fly *Drosophila melanogaster*, no evidence for gBGC has been reported, albeit a weak effect on the X chromosome (Galtier, Bazin, et al., 2006; MC Robinson et al., 2014). In contrast, Wall-berg et al. (2015) estimated the genome average *B* to be above 5 in the honey bee *Apis mellifera*, again with substantial variation between low-recombining and high-recombining regions. Note that *N*_*e*_ is expected to be much smaller in *Homo sapiens* and the eusocial *A. mellifera* than in *D. melanogaster* (Romiguier, Lourenco, et al., 2014). Galtier, Roux, et al. (2018) analysed site frequency spectrum at synonymous positions in the coding sequences of 30 species of animals. They estimated that the average *B* at third codon positions varies between 0 and 2 among species, without any significant relationship with *N*_*e*_-related life history traits. These comparisons among distantly-related animals revealed substantial variation in the intensity of gBGC among species, but, somewhat paradoxically, no detectable effect of *N*_*e*_.

Another attempt to quantify the strength of gBGC is to get information from between-species divergence data, instead of within-species polymorphism data. Capra et al. (2013) simultaneously modelled the effects of purifying selection and gBGC during the divergence between humans and chimpanzees and estimated that in apes 0.33% of the genome is undergoing gBGC at rate *B* = 3. This is lower than the estimates provided by Glémin et al. (2015, see above), presumably because Capra et al. (2013) assumed a constant gBGC rate at any location of the genome, whereas recombination hotspots are known to be highly dynamic in apes (Auton et al., 2012; Lesecque et al., 2014). Using a method that combines polymorphism and divergence data, De Maio et al. (2013) estimated the average *B* to be of the order of 0.3-0.7 in apes, consistent with Glémin et al. (2015). Lartillot (2013) analysed coding sequence divergence in 33 species of placental mammals and estimated the among-gene and among-species variation of *B*. He found that the average *B* varied among species from *∼* 0.1 (in apes) to 3-5 (in bats and lagomorphs), with an among-gene standard deviation of *B* as high as twice the mean. Lartillot (2013) detected a significant, negative correlation between *B* and species body mass. Body mass being strongly and negatively correlated with population density in mammals (Damuth, 2007), this result suggests that *N*_*e*_ might be a determinant of the strength of gBGC in mammals, in agreement with theoretical expectations. Elaborating on the approach of De Maio et al. (2013), Borges et al. (2019) also reported a positive relationship between the population scaled gBGC coefficient and *N*_*e*_ across species/populations of apes.

So on one hand comparative analyses of site-frequency spectra among animals did not reveal any effect of *N*_*e*_ on the strength of gBGC, while on the other hand the analysis of the substitution pattern in mammals is consistent with a *N*_*e*_ effect. Also a bit surprisingly, the estimated range of variation of *B* across mammals (0.1-5, Lartillot, 2013) is wide enough to contain all the estimates of *B* reported in any species of animals so far. Lartillot (2013) analysed a subset of currently annotated mammalian orthologs (1329 exons in the largest data set), and importantly, relatively ancient divergences, at the family or order level, thus capturing the average effect of gBGC across dozens of million years. Here we analyse a large set of genes from closely related species in four families of mammals, thus accessing a high-resolution map of the effect of gBGC on coding sequences, both in time and across the genome. We focus on two key features of gBGC-driven molecular evolution, namely clustered AT*→*GC substitutions, and an excess of AT*→*GC over GC*→*AT substitutions compared to the mutation process. Estimating *B* in 40 lineages of mammals, we show that gBGC explains a substantial fraction of synonymous and non-synonymous AT*→*GC substitutions, that *N*_*e*_ is a strong predictor of the intensity of gBGC in mammals, and that large-*N*_*e*_ and small-*N*_*e*_ taxa differ substantially in how gBGC is distributed among and within genes.

## Results

### Overview

We analysed patterns of AT*→*GC (i.e., Weak to Strong, or WS), GC*→*AT (SW) and GC-conservative (SSWW) coding sequence nucleotide substitutions in 40 recently diverged lineages (branches) from four families of mammals (Fig.1), namely Hominidae (humans and apes), Cercopithecidae (old world monkeys), Bovidae (cattle, sheep and allies) and Muridae (mice, rats, gerbils). A total of 1,104,917 third codon position synonymous substitutions and 514,552 first or second codon position non-synonymous substitutions were called. The median number of substitutions across branches was 24,960, and the minimum was 3927. The overall ratio of non-synonymous to synonymous substitutions, *d*_*N*_*/d*_*S*_, was 0.233; the family-specific *d*_*N*_*/d*_*S*_ ratio was 0.275, 0.252, 0.228 and 0.213 in Hominidae, Cercopithecidae, Bovidae and Muridae, respectively. The *d*_*N*_*/d*_*S*_ ratio is a marker of *N*_*e*_ in mammals, with small populations experiencing a higher substitution load, hence a higher *d*_*N*_*/d*_*S*_ (Nikolaev et al., 2007; Popadin et al., 2007; Romiguier, Figuet, et al., 2012). These results therefore indicate that the four families of our data set rank in the Hominidae *<* Cercopithecidae *<* Bovidae *<* Muridae order as far as *N*_*e*_ is concerned, consistent with previous analyses (Lartillot, 2013; Romiguier, Ranwez, et al., 2013).

**Figure 1.**
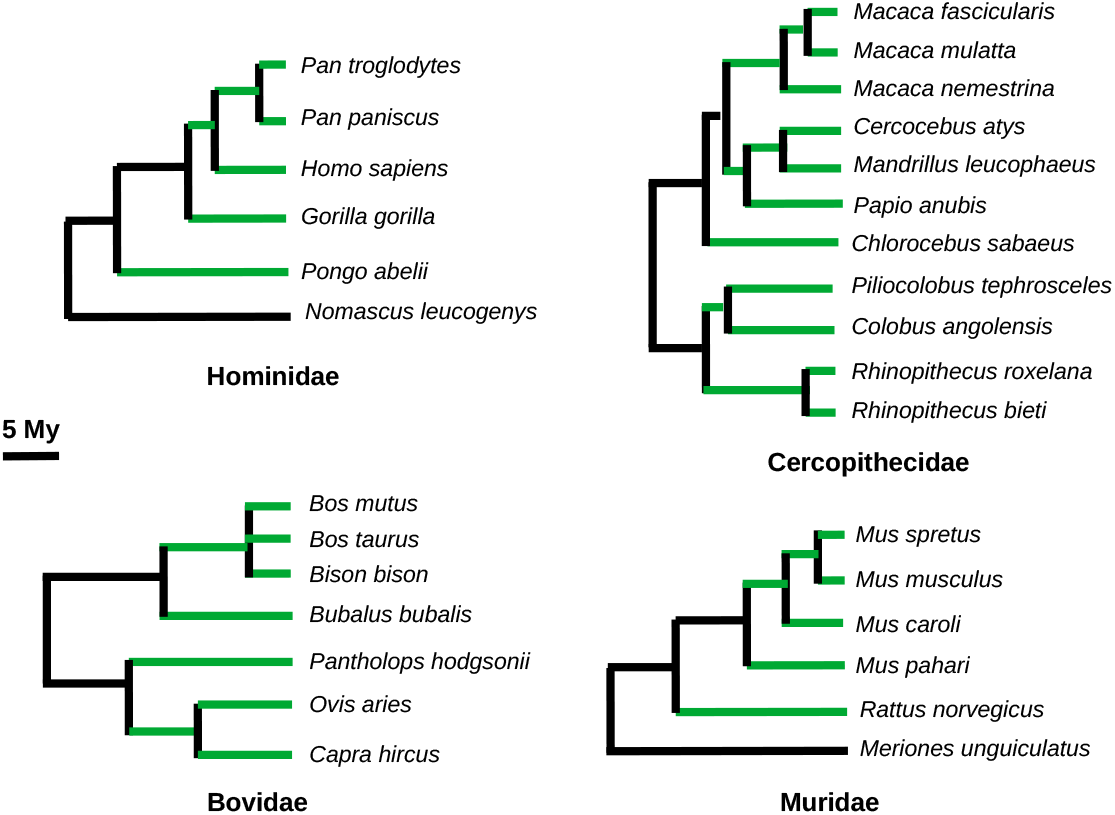
The four families, 32 species and 40 lineages of mammals (green branches) analysed here. Branch lengths are proportional to the estimated divergence time.

### Substitution clustering

Focusing on synonymous substitutions, we calculated Moran’s *I* (Moran, 1950), a statistics that measures spatial aurocorrelation and was adjusted to target the 400 bp scale. This index therefore measures the tendency for substitutions (of a specific sort) having appeared in a given branch to be located less than 400 bp apart. Fig.2 shows the distribution among branches of the average centered Moran’s *I*, separately for WS and SW synonymous substitutions. The centered Moran’s *I* for SW substitutions was very close to zero in all branches from all four families, indicating very little, if any, clustering of substitutions. WS substitutions behaved differently: the centered Moran’s *I* was close to zero in Hominidae, perceptibly positive in Cercopithecidae, and reached much higher values in Bovidae and Muridae, demonstrating the existence of clusters of synonymous WS substitutions in these two families. This pattern - clustering of WS but not SW substitutions - is a signature of gBGC (*e*.*g*. Dreszer et al., 2007); its intensity appears to increase with *N*_*e*_ across the four families analysed here.

**Figure 2.**
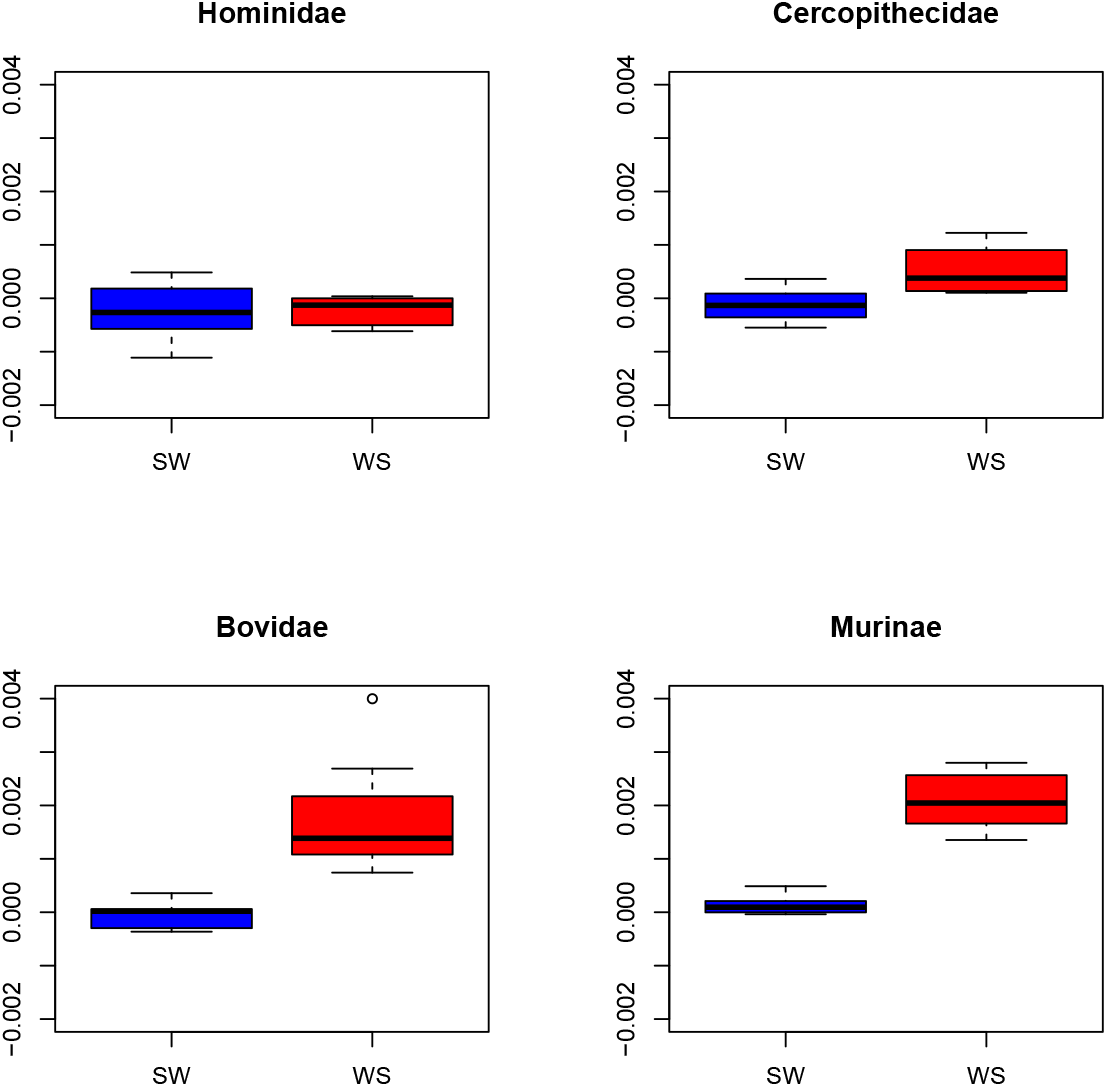
Distribution of centered Moran’s *I* for WS and SW synonymous substitutions in four families of mammals. Only branches in which at least 100 genes had at least 3 inferred substitutions were included.

Simulations were performed in order to assess the amount of clustering needed to explain the observed values of Moran’s *I*. Our simulation procedure considers two levels of clustering, one at the 500 bp scale and one at the 40 bp scale, while accounting for the intron-exon structure and the among-genes variance in mutation rate and GC-content (see Methods). In Muridae and Bovidae, we were able to replicate the observed values of Moran’s *I* when 15-40% of the simulated substitutions appeared in clusters. This percentage was 0-10% in Cercopithecidae, and non-existent in Hominidae (Supplementary Fig. S1).

### Estimating *B*

For each branch we estimated the population gBGC coefficient *B* = 4*N*_*e*_*b* and its variation based on synonymous WS, SW and SSWW synonymous substitution counts. Various models were fitted to the data via the maximum likelihood (ML) method, assuming that the mutation process is known (TCA Smith et al., 2018). Model M1 assumes a constant intensity of gBGC, *B*, among and within genes. Model M2 considers two categories of genes, each with its own gBGC intensity, assumed to be shared by all sites within a gene. M2 led to a rejection of M1 by a likelihood ratio test (*p* − *val <* 0.05) in 36 branches out of 40. The M3z model assumes three categories of genes, which we below denote “cold” (*B* = 0), “mild”, and “hot”. M3z rejected M1 in 39 branches out of 40, and M2 in 27 branches. There was, therefore, strong evidence for a variable *B* across genes in this data set.

Then we fitted models that assume some variation of *B* both among and within genes. Model M3h considers three categories of genes that differ in terms of the prevalence, *q*, of gBGC hotspots. gBGC is assumed to operate at intensity *B*_*h*_ within hotspots, and zero outside hotspots. Model M3sh is a simplified version of M3h obtained when *q* approaches zero. Applying these two models led to a dramatic increase in log-likelihood for most branches (Supplementary Table S1), which is indicative of the existence of substantial within-gene variation in gBGC intensity. Model M3h rejected M3sh by a likelihood ratio test only in one branch out of 40 (*Bison bison* terminal branch, Bovidae), consistent with the idea that gBGC hotspots occupy a small fraction of coding sequence length.

The across-genes average gBGC intensity, 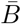, varied among models, with models allowing for more variation in *B* usually yielding a higher 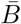 (Supplementary Table S1). Below we report estimates of 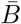 obtained under the M3sh model. These were very similar to estimates obtained by averaging 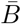 across the M1, M2, M3z, M3sh and M3h models, weighting by the AIC of each model (Posada and Buckley, 2004, Supplementary Fig. S2).

Fig.3 shows the distribution of 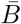 among branches in the four analysed families. The median 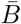 was just below 0.5 in primates, 0.82 in Bovidae and 1.76 in Muridae. We calculated the across genes relative standard deviation (RSD) of *B*, which is the ratio of the standard deviation by the average *B*. The RSD would be expected to be constant across branches if the across-genes distribution of the intensity of gBGC only differed among branches by a coefficient of proportionality. We found that the RSD was generally rather high (median RSD across branches: 1.8), and substantially smaller in Muridae (median: 1.2) than in the other three families (median Bovidae: 1.8; median Hominidae: 1.7; median Cercopithecidae: 2.1). This suggests that the intensity of gBGC is more evenly distributed among genes in Muridae than in the other taxa. Of note, this result superficially appears to contradict the analysis illustrated by Fig.2, which shows that the clustering of WS substitutions is maximal in Muridae. Importantly, the Moran’s *I* analysis (Fig.2) addresses the within-gene clustering of substitutions, whereas in the RSD analysis we consider the among-gene variation in gBGC intensity.

**Figure 3.**
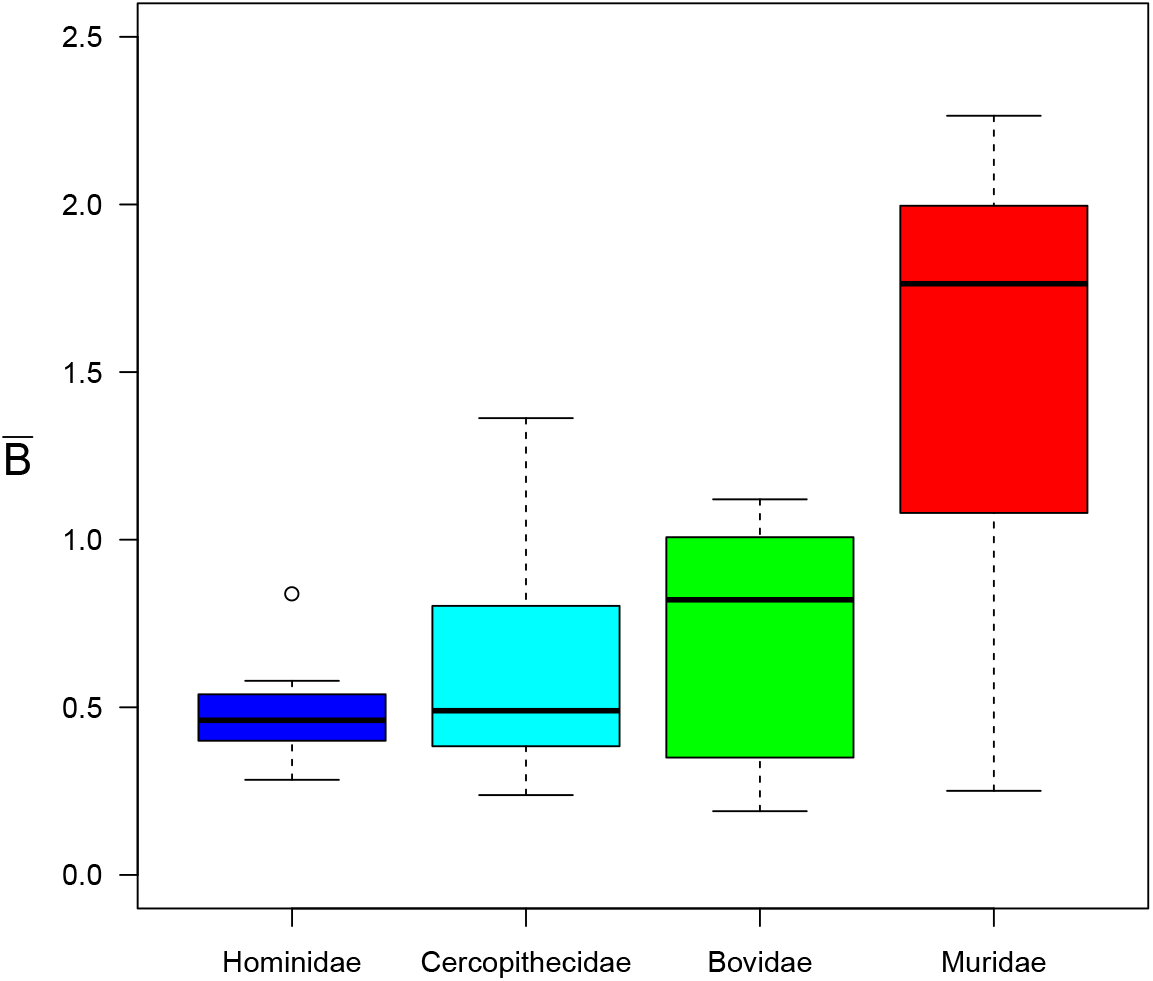
Distribution of the average estimated *B* (M3sh model). One outlying data point is missing from the figure: the estimated average *B* was 3.86 in the (*Capra hircus, Ovis aries*) ancestral branch (Bovidae).

We estimated in each branch the number of WS substitutions that would be expected in the absence of gBGC. This was achieved by forcing *B* = 0 for all categories of genes under the M3sh model (see Methods). We found that gBGC results in a substantial excess of WS substitutions, which varies from typically 30% in primates to typically 60% in Muridae (Supplementary Fig. S3). No effect of gBGC on SW substitutions is expected under the M3sh model (see Methods).

### 0.1 Correlates of *B*

We correlated the log-transformed estimated 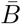 with log-transformed branch-specific *d*_*N*_*/d*_*S*_ ratio and found a significantly negative relationship (*n* = 40; *r*^2^ = 0.24; p-val=0.0013). The correlation coefficient of the 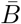 vs. *d*_*N*_*/d*_*S*_ relationship was also significantly negative when calculated within Hominidae (*n* = 7; *r*^2^ = 0.83; p-val=0.0043), within Bovidae (*n* = 9; *r*^2^ = 0.79; p-val=0.0013) and within Muridae (*n* = 7; *r*^2^ = 0.57; p-val=0.049). No significant relationship was detected within Cercopithecidae (Fig.4, left). Very similar results were obtained when we correlated the estimated 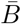 with *d*_*N*_*/d*_*S*_ calculated based on SSWW substitutions only, i.e., a statistics essentially independent of gBGC: the squared correlation coefficients were 0.82 (p-val=0.0047), 0.68 (p-val=0.0059) and 0.73 (p-val=0.0139) within Hominidae, Bovidae and Muridae, respectively, and 0.23 (p-val=0.0017) for the whole data set (all variables log-transformed). A literature search yielded estimates of heterozygosity (*i*.*e*., within-species genetic diversity), *π*, in 18 species of our data set. The estimated 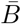 was positively correlated with *π* (*r*^2^ = 0.73; p-val=2.1 *×* 10^−5^; fig.4, right). The sample size was here too small to investigate the within-family relationships. B was also found to be negatively correlated with species longevity (*r*^2^ = 0.36, p-val=0.0026) and log-transformed body mass (*r*^2^ = 0.22, p-val=0.017).

**Figure 4.**
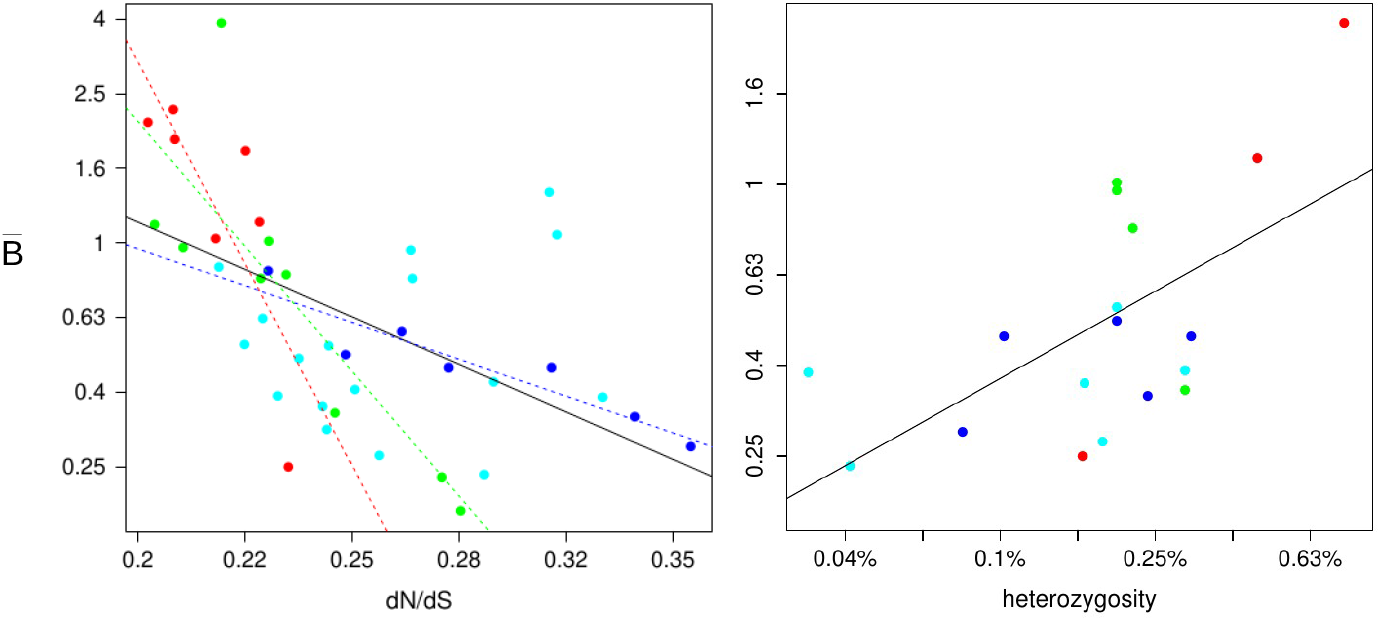
Relationship between the average estimated *B* and *dN/dS* (left panel, *n* = 40 branches) or heterozygosity (right panel, *n* = 18 species) across 40 mammalian lineages in log-transformed scales. *B* was estimated under the M3sh model. Blue: Hominidae; cyan: Cercopithecidae; green: Bovidae; red: Muridae; black line: regression line for the whole data set; colored dotted lines: family-specific regression lines

### Substitution clustering conditional on *B*

Fig.2 revealed virtually no clustering of WS substitutions in Hominidae, even though the analysis of substitution counts demonstrated a significant impact of gBGC on coding sequences in this family (Fig.3). To test whether the spatial distribution of WS substitutions really differs between mammalian families, we analysed substitution clustering conditional on *B*. For each branch, we first fitted to WS, SW and SSWW substitution counts a gBGC model, M5f, assuming five categories of genes undergoing distinct gBGC intensities, from *B* = 0 in the coldest category to *B* = 10 in the hottest one. We assigned each gene to one of these gBGC intensity categories, and calculated the average Moran’s *I* for WS substitutions separately for the five categories (Fig.5). This was also done using the hotspot version of this five-category model, M5shf (Supplementary Fig. S4). The size of dots in Fig.5 and Supplementary Fig. S4 reflects the proportions of the five classes of genes in each family, genes from distinct branches being here merged. We found that the average Moran’s *I* increased with gBGC intensity, as expected, but varied strongly among families in every gBGC category, with Muridae consistently showing the highest average Moran’s *I*, and Hominidae the lowest, at all gBGC intensities. This result indicates that the level of clustering of WS substitutions differs across families to an extant that cannot be explained just by differences in average *B*.

**Figure 5.**
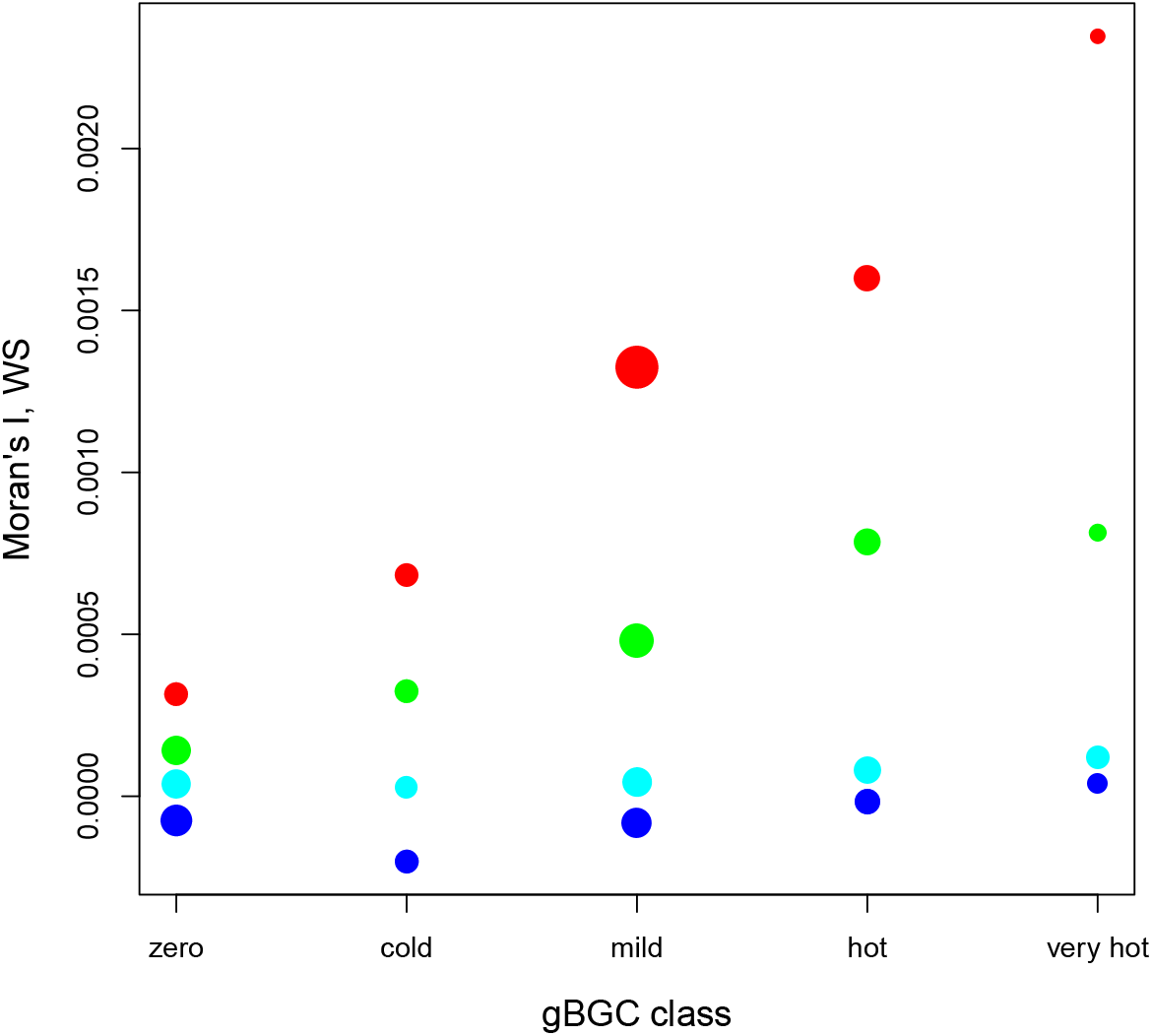
Average centered Moran’s *I* as a function of average estimated gBGC strength *B*. Each dot is for a class of genes in a family (all species merged). Genes were assigned to classes under the M5f model. Dot size reflects the relative number of genes in the considered class. Blue: Hominidae; cyan: Cercopithecidae; green: Bovidae; red: Muridae.

## Discussion

Analysing the substitution pattern in coding sequences across four mammalian families, we checked two predictions of the gBGC model, namely a clustering of WS substitutions and an excess of WS over SW substitutions compared to the mutation pattern. Both approaches revealed a conspicuous effect of gBGC in mammalian coding sequences.

### Ubiquitous gBGC in mammals

Dreszer et al. (2007) investigated the substitution pattern in the human genome and showed that clusters or nearby substitutions tend to be enriched in the WS sort. The effect, although significant, was not particularly strong: the proportion of WS substitutions in clusters reached 0.55, whereas it was 0.44 on average (their figure 1A). Analyzing exon evolution in apes, Berglund et al. (2009) and Galtier, Duret, et al. (2009) identified a few dozens of GC-biased exons, out of 10,000 analyzed exons. Here we applied a distinct but related approach to mammalian coding sequences, and reveal only a weak, if any, tendency for WS synonymous substitutions to be clustered in Hominidae, consistent with previous research. The trend, however, was obvious in Cercopithecidae, and strong in Bovidae and Muridae (Fig.2). Our simulations suggest that 15%, and maybe up to 40%, of WS synonymous substitutions appear as clusters in these two families. These WS substitution clusters likely reflect a localised effect of gBGC at recombination hotspots. Here we show that such clusters, although anecdotical in humans and apes, are a major component of the substitution pattern in other families of mammals.

Our estimate of the *B* parameter, which measures the average intensity of gBGC across genes, varied between 0.2 and 3.9 among the 40 analysed lineages. In primates, the median estimated 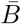 was *∼* 0.5, i.e., in the range of previously published values: 0.1 in hominoids (Lartillot, 2013), 0.38 in humans (Glémin et al., 2015), 0.35-0.7 in apes (De Maio et al., 2013). Our estimates of 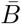. in Bovidae (*∼* 0.5 − 1) and Muridae (*∼* 1 − 2) are also quite similar to those obtained by Lartillot (2013) in the *Bos taurus* (Bovidae), *Mus musculus* and *Rattus rattus* (Muridae) lineages. In Bovidae, we found a positive relationship between the estimated 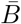 and branch age (in million years), defined as the average between the date of the top and bottom nodes of a branch (*n* = 9 branches; *r*^2^ = 0.75; p-val=0.003). This is consistent with the hypothesis of a high ancestral *N*_*e*_ in this taxon, as also suggested by fossil data and *d*_*N*_*/d*_*S*_-based reconstructions (Figuet, Ballenghien, Lartillot, et al., 2017; Figuet, Romiguier, et al., 2014). Our study could not confirm the report by Romiguier, Ranwez, et al. (2010) and Lartillot (2013) of a particularly strong gBGC in bats, tenrecs and lagomorphs due to the unavailability of fully-sequenced, closely related species in sufficient numbers in these taxa.

The estimated genome average *B* was in the nearly neutral zone in the four families analysed here. Even so, gBGC was found to be pervasive and strongly impact the substitution process in coding sequences. Fitting a three-category, hotspot model across genes, we estimate that 30 to 60% of the WS synonymous substitutions can be attributed to gBGC in mammals. It should be noted that this estimate, as well as the estimates of *B* we report in this work, is dependent on the assumption of a known and constant mutation process. Here we used the WS, SW, SS and WW mutations rates obtained from TCA Smith et al. (2018), who analysed 130,000 *de novo* mutations inferred from mother/father/child trios in humans - a very large data set. Milholland et al. (2017) compared the germline mutation pattern of *H. sapiens* and *M. musculus* and did not detect any conspicuous difference between the two species in the proportions of SW, WS and SSWW mutations, and neither did Wang et al. (2020) when comparing *Macaca mulatta* (Cercopithecidae) to *H. sapiens*. So the existing literature does not seem to question our assumption of constant relative mutation rates in mammals - but note that no such data is available in Bovidae, to our knowledge. Also note that the clustering analysis (Fig.2) does not make any assumption regarding the mutation process.

It should be noted that our estimate of *B* in this analysis is based on substitution counts inferred via a parsimony-based approach. This way of counting substitutions is not devoid of potential problems. Maximum parsimony substitution inference is known to be biased towards common-to-rare changes (Eyre-Walker, 1998). However, the relatively recent divergence times we are considering presumably keeps this effect to a minimum. Indeed the longest branch across all four trees (Fig. 1), the *Rattus norvegicus* terminal branch, has a length below 0.08 substitutions per site, which is the minimal length for which this problem was detectable in Eyre-Walker, 1998. Using closely related species, on the other hand, runs into another potential bias: when closely related species are analysed, there is a risk that within-species polymorphism contributes a non-negligible fraction of the observed sequence variation, biasing the estimation of quantities such as the *d*_*N*_*/d*_*S*_ ratio (Mugal, Kutschera, et al., 2020). This bias likely affects the estimation of the relative SW and WS substitution rates as well, since the expected SW/WS rate ratio differs between polymorphism and divergence when gBGC is at work. More work would be needed to confirm and quantify the effect of this bias on our analysis.

### A significant effect of the effective population size

Although a significant effect of gBGC was detected in all four analysed families, its intensity varied conspicuously among families, Muridae being the most strongly impacted, followed by Bovidae, Cercopithecidae, and Hominidae. It is noticeable that gBGC ranks family in the same order as *N*_*e*_, as measured by the family-average *d*_*N*_*/d*_*S*_ ratio, this order being consistently recovered in nearly all the analyses we performed. This is in line with the expectation that the intensity of gBGC should be higher in large than in small populations. An effect of *N*_*e*_ was also detected by correlating 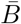. with the *d*_*N*_*/d*_*S*_ ratio across branches, and with heterozygosity across species (Fig.4). These analyses confirmed the significance of the effect, both among and within families, thus corroborating the relationship uncovered by Lartillot (2013) at a deeper phylogenetic scale.

Interestingly, the slope of the 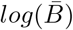 vs. *log*(*d*_*N*_*/d*_*S*_) relationship differed conspicuously among families in Fig.4. This result might tell something about the strength of selection on amino-acid changing mutations in mammalian coding sequences. Welch et al. (2008) showed that, assuming a Gamma distribution of deleterious effects of non-synonymous mutations, the *d*_*N*_*/d*_*S*_ ratio is expected to be proportional to *N*_*e*_^−*β*^, where *β* is the shape parameter of the Gamma distribution. So under this assumption, and since *B* is proportional to *N*_*e*_, the slope of the 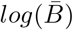 vs. *log*(*d*_*N*_*/d*_*S*_) relationship should equal −1*/β*. This rationale yields estimates of *β* equal to 0.51 in Hominidae, 0.15 in Bovidae, and 0.09 in Muridae. These figures differ considerably from estimates obtained by site frequency spectrum analyses, *i*.*e*., *β ∼* 0.15 in Hominidae and *∼* 0.2 in Muridae (Castellano et al., 2019; Galtier and Rousselle, 2020; Huber et al., 2017). More work is needed to understand the origin and meaning of this discrepancy. At any rate, our results suggest that gBGC analysis could constitute a new source of information on the variation in *N*_*e*_ among species, and might enrich the ongoing discussion on this issue (e.g Buffalo, 2021; Galtier and Rousselle, 2020).

The among lineages correlation between the estimated 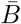 and the *d*_*N*_*/d*_*S*_ ratio we report here in mammals, which confirms Lartillot (2013)’s results, contrasts with the absence of such a correlation at the Metazoa scale. Large-*N*_*e*_ fruit flies and marine molluscs, for instance, are less strongly impacted by gBGC than small-*N*_*e*_ bees and amniotes (Galtier, Roux, et al., 2018; MC Robinson et al., 2014; Wallberg et al., 2015). The simplest explanation for this is that *b*, the transmission bias, probably differs much between distantly related taxa, due to differences in recombination rate, repair bias and/or conversion tract length. For instance, the recombination rate is known to be particularly high in honey bees (Wilfert et al., 2007).

Two recent studies experimentally assessed the intensity of gBGC in mice via crosses followed by sperm (Gautier, 2019) or progeny (Li et al., 2019) whole genome sequencing. Both estimated that *b* is lower in mice than in humans - maybe five times lower, although this figure requires confirmation. Gautier (2019) invoked purifying selection against gBGC to explain this result. Indeed, because of its deleterious effects (Berglund et al., 2009; Galtier, Duret, et al., 2009; Necşulea et al., 2011), gBGC as a process could be counter-selected, and purifying selection being more effective in large than in small population, this verbal model would predict a lower *b* in mice than in human. Adapted to our results, this hypothesis of a negative correlation between *N*_*e*_ and *b* would imply that the range of variation in *B* should be narrower than the range of variation in *N*_*e*_ among mammalian lineages. We indeed observed a narrower variation in the magnitude of the estimated 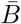 (standard deviation of 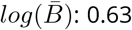) than of heterozygosity (standard deviation of *log*(*π*): 0.74; same 18 species used in these two calculations, see Fig.4, right panel). The difference, however, is not particularly pronounced, and does not suggest the existence of a strong negative relationship between *b* and *N*_*e*_ in mammals. That said, not only *N*_*e*_ influences the variation of *π*: the mutation rate also matters. Among-species differences in per generation mutation rate, if any, should be taken into account for a better assessment of the *b* vs. *N*_*e*_ relationship (Brevet and Lartillot, 2021).

### Recombination hotspots dynamics

The among-gene variation in gBGC intensity, measured by the relative standard deviation of *B*, was found to be substantial in all branches, while lower in Muridae than in the other three families - a pattern also reported by Lartillot (2013). gBGC seems to be more evenly distributed across the genome in this taxon, consistent with previous reports that GC3 in murid rodents has been increasing and was homogenised since the common ancestor of this family (Clément and Arndt, 2011; Mouchiroud et al., 1988; M Robinson et al., 1997; Romiguier, Ranwez, et al., 2010). Muridae appears to be a peculiar group of mammals with this respect (Romiguier, Ranwez, et al., 2010), and one should keep this in mind when interpreting patterns of gBGC-related evolution in this taxon. Of note, the existing literature does not suggest that the recombination map is less heterogeneous in mouse or rat than in primates (Brunschwig et al., 2012; Jensen-Seaman et al., 2004; McVean et al., 2004).

The within-gene heterogeneity in *B*, in contrast, was more pronounced in Muridae than in Bovidae and, particularly, primates (Fig.2), and this was true even when we controlled for gene-specific *B* (Fig.5): for any particular intensity of gBGC, WS substitutions tend to be more clustered in large-*N*_*e*_ than in small-*N*_*e*_ species. This intriguing result might be interpreted in relation with the dynamics of recombination hotspots. To result in a cluster of WS substitutions, a recombination hotspot must be active during a sufficiently long period of time for several WS alleles to reach a high population frequency. We suggest that in primates gBGC does not generate a pattern of highly clustered WS substitutions because heterozygosity is low and recombination hotspots are short-lived in this taxon. Indeed, recombination hotspots are known to be particularly ephemeral in Hominidae, due to a transmission distortion associated with the Red Queen-like evolution of the major hotspot determining gene PRDM9 (Auton et al., 2012; Coop and SR Myers, 2007; S Myers et al., 2010). For instance, Lesecque et al. (2014) showed that denisovians and modern humans did not share the same recombination hotpots, while the level of divergence between these two genomes is of the order of one synonymous substitution per gene on average (Meyer et al., 2012). Our results might suggest that the situation differs in other taxa of mammals, maybe in a way related to *N*_*e*_. At any rate, heterozygosity is higher in large-*N*_*e*_ species, which increases the probability that a given local episode of gBGC results in more than one WS substitutions, irrespective of recombination hotspot lifespan. A deeper understanding of this result would require to account for the many factors influencing the turnover time of PRDM9 alleles and recombination hotspots (Latrille et al., 2017), the length of gene conversion tracts, as well as the population mutation rate and fixation probability of WS mutations.

### Concluding remarks

Quantifying gBGC in closely related species of mammals, we report a pervasive effect on the nucleotide substitution process, a positive relationship with *N*_*e*_, and a complex pattern of variation within and among genes. This work also demonstrates that the analysis of gBGC has the potential to illuminate various aspects of molecular evolution, including the distribution of fitness effect of mutations and the dynamics of recombination hotspots. The apparent lack of a *N*_*e*_ effect on gBGC intensity at the Metazoa scale is an unresolved question that requires further quantification of the strength of gBGC in non-vertebrate taxa.

## Material and Methods

### Sequence data

Mammalian coding sequence alignments were downloaded from the Orthomam v10 database (Scornavacca et al., 2019). The four families of mammals represented by at least six species in OrthoMam v10 were selected, namely Hominidae (six species, 11,859 genes), Cercopithecidae (eleven species, 10,834 genes), Bovidae (seven species, 9527 genes), and Muridae (six species, 11,758 genes). In Bovidae, the *Bos indicus* sequences were not considered since this taxon is a subspecies of *Bos taurus*. In all four families the phylogenetic histories of the sampled species are well documented, with the exception of the unresolved relationship between cattle, yak and bison (Fabre, Hautier, et al., 2012; Fabre, Rodrigues, et al., 2009; Hassanin et al., 2012; Vanderpool et al., 2020, Fig1). Nodes were dated based on the TimeTree website (http://www.timetree.org/) using the median date estimates.

### Substitution mapping

For each of the four data sets, nucleotide substitutions were mapped to the resolved branches of the trees using a stringent parsimony approach. For any given branch, an X*→*Y substitution was recorded if and only if all species descending from the considered branch carried state Y, and all other species carried state X. All positions not matching this exact pattern, including positions with missing data or gaps, were disregarded. Branches connected to the root of the tree were excluded, as well as branches whose number of descending species was higher than half the total number of sampled species in the family. A total of 40 distinct branches were considered - seven in Hominidae and Muridae, nine in Bovidae, 17 in Cercopithecidae (Fig.1). For each branch and each coding sequence, the number and positions of non-synonymous and synonymous substitutions were recorded, distinguishing the AT*→*GC (WS), GC*→*AT (SW) and GC-conservative (SSWW) sorts. Only synonymous substitutions occurring at third codon positions, and non-synonymous substitutions occurring at first or second codon positions, were counted. For any given branch, substitutions that mapped to consecutive sites were ignored, and genes in which the per base pair substitution rate was higher than ten times the across-genes median rate were discarded (implying that the number of analysed genes could slightly differ among lineages within a family). The last two steps aimed at diminishing the effect of misaligned regions.

### Clustering analysis (synonymous substitutions)

For each branch and each gene of length above 400 bp, we calculated Moran’s *I* index (Moran 1950) separately for WS and SW synonymous substitutions. We used a weight matrix defined as follows: the weight equalled one for any two substitutions distant of 400 bp or less, and zero for any two substitutions more distant than 400 bp. Window widths of 200 bp and 100 bp gave qualitatively similar results. For each branch and each sort of substitutions, Moran’s *I* was averaged across genes, excluding genes with less than three substitutions of the considered sort. Moran’s *I* has a negative expectation of −1*/*(*l* − 1) under the null hypothesis of no spatial autocorrelation, where *l* is the number of third codon positions of the considered gene. Here we used the centered version of the statistics, *I* + 1*/*(*l* − 1), the expectation of which is zero in the absence of substitution clustering.

### Clustering: simulations

We downloaded from the Ensembl database coding sequence annotations in one representative species per family, namely *Homo sapiens* (Hominidae), *Macaca mulatta* (Cercopithecidae), *Bos taurus* (Bovidae) and *Mus musculus* (Muridae), disregarding coding sequences shorter than 400 bp. Then we simulated substitution data in a hypothetical branch by iteratively sampling the location of third-codon-position substitutions across coding sequences using the following method:

(initiation:) randomly sample the location of the first substitution among the third codon positions of all genes;

(iteration:)

- with probability 1 − *p*_*clust*_, randomly sample the location of the (*n* +1)^*th*^ substitution among the third codon positions of all genes;
- with probability *p*_*clust*_, randomly sample the location of the (*n* + 1)^*th*^ substitution in the neighborhood of the *n*^*th*^ substitution (clustered substitutions). More precisely, conditional on the *n*^*th*^ and (*n* + 1)^*th*^ substitutions being clustered,
- with probability *p*_*CO*_ the (*n* + 1)^*th*^ substitution was randomly sampled in a window of width *l*_*CO*_ centered on the location of the *n*^*th*^ substitution, and
- with probability *p*_*NCO*_ = 1 − *p*_*CO*_ the (*n* + 1)^*th*^ substitution was randomly sampled in a window of width *l*_*NCO*_ centered on the location of the *n*^*th*^ substitution.

This was intended to represent the fact that gene conversion tracts associated to crossing-over and non-crossing-over events are of different lengths (Cole, Baudat, et al., 2014). If the sampled location of the (*n* + 1)^*th*^ substitution reached beyond the boundaries of the exon carrying the *n*^*th*^ substitution, then the (*n* + 1)^*th*^ substitution was ignored. Our procedure also accounted for the existence of variation in mutation rate among exons: we assumed that one half of the exons had a mutation rate *γ* times as high as the other half. We separately simulated WS and SW substitution data, accounting for the distribution of GC-content at third codon positions - hence, the availability of W and S sites - in the four groups. A gene with GC3=90% was 10 times more likely to host a SW substitution than a gene with GC3=10% in our simulations.

Two parameters of the simulation procedure were varied among conditions, namely the per third codon position density of substitutions (taking values in {0.0003, 0.001, 0.003, 0.01, 0.03}) and the probability *p*_*clust*_ for two successive substitutions to be clustered (taking values in {0, 0.1, 0.2, 0.3, 0.4}). The other parameters were fixed to constant values estimated from the literature. Parameters *l*_*CO*_ and *l*_*NCO*_ were set to 500 and 40 bp, respectively (Cole, Baudat, et al., 2014; Jeffreys and May, 2004; Li et al., 2019; Williams et al., 2015). Parameter *γ* was set to 3, ensuring an among-exon mutation rate relative standard deviation of 0.5, in agreement with figure 4 in TCA Smith et al. (2018). Finally, parameter *p*_*CO*_ was set to 0.86, according to the following rationale: the CO/NCO odds ratio for the first substitution to occur in a gene conversion tracts is (*l*_*CO*_*n*_*CO*_)*/*(*l*_*NCO*_*n*_*NCO*_), where *n*_*CO*_ and *n*_*NCO*_ are the number of crossing-over and non crossing-over events, respectively; the CO/NCO odds ratio for the second substitution to occur in the same gene conversion tract is *l*_*CO*_*/l*_*NCO*_; so the CO/NCO odds ratio for the occurrence of a pair of clustered substitutions is the product, *p*, of the two terms above, and *p*_*CO*_ = *p/*(1 + *p*). Using *n*_*CO*_*/n*_*NCO*_ = 0.1 (Baudat and Massy, 2007; Cole, Kauppi, et al., 2012), *l*_*CO*_ = 500 and *l*_*NCO*_ = 40 we obtain *p*_*CO*_ = 0.86.

### Maximum likelihood estimation of gBGC strength

For each branch and each gene, we counted the numbers of inferred WS, SW and SSWW synonymous substitutions at third codon positions. Then we fitted mutation/gBGC/drift models to these observations in the maximum likelihood framework.

Consider a coding sequence of length *l* evolving in a panmictic diploid population of constant size *N*_*e*_ under neutrality during a period *T* of time. The expected number of substitutions, *n*^*^, depends on the mutation rate *μ* and fixation probability *f* :

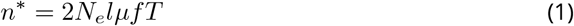

Assuming a homogeneous gBGC intensity of *b*, the fixation probability of WS, SW and SSWW mutations can be written as:

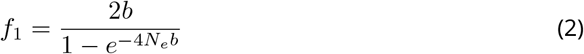

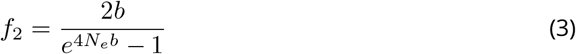

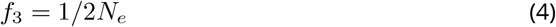

Here and below, subscript 1, 2 and 3 respectively refer to the WS, SW and SSWW sorts of change. Substituting in equation 1 and only considering third codon positions, we get the expected number of synonymous substitutions of the three sorts:

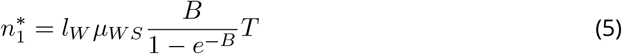

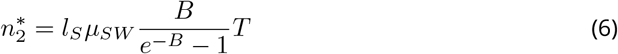

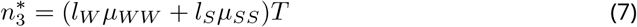

where *B* = 4*N*_*e*_*b, l*_*W*_ and *l*_*S*_ are the number of AT- and GC-ending codons, respectively, in the considered coding sequence, and *μ*_*WS*_, *μ*_*SW*_, *μ*_*SS*_ and *μ*_*WW*_ are the corresponding mutation rates.

Assuming that the number of WS substitutions is Poisson distributed, the probability of observing *n* WS substitutions given *B* and *T* is given by the following function:

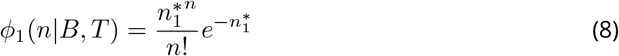

and similarly for SW and SSWW substitutions:

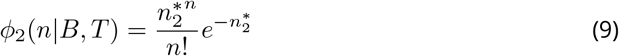

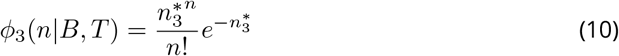

We modelled the variation of *B* among genes using discrete distributions. Assume there are *k* categories of genes, with each category including a fraction *p*_*k*_ of the genes and characterised by a population-scaled gBGC intensity *B*_*k*_, assumed to be constant within genes. For a gene at which *n*_1_, *n*_2_, and *n*_3_ substitutions of the three sorts are observed, the likelihood can be written as:

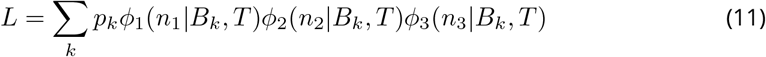

We considered various models that differ in how *B* varies across genes. Under model M1, a constant *B* across genes is assumed. Model M2 defines two categories of genes, each with its own gBGC intensity. Model M3z (for zero) has three categories of genes, among which one has a gBGC intensity of zero, the other two being free parameters. Model M5f (for fixed) has five categories of genes with fixed gBGC intensities equal to *B*_1_ = 0, *B*_2_ = 0.333, *B*_3_ = 1, *B*_4_ = 3.333 and *B*_5_ = 10, respectively. In all four models the proportions of genes in the various categories were free to vary, *T* was assumed to be shared among genes and the relative mutation rates *μ*_*WS*_, *μ*_*SW*_, *μ*_*SS*_ and *μ*_*WW*_ were set to empirical estimates obtained from Smith et al. (2018), i.e., *μ*_*WS*_ = 5.21, *μ*_*SW*_ = 10.90, *μ*_*SS*_ = 2.07 and *μ*_*WW*_ = 1.

The models above assume a homogeneous rate of gBGC among positions within a gene. To account for the existence of hotspots of gBGC, we modelled the within-gene variation of *B* by assuming that only a fraction *q* of the positions undergo gBGC at rate *B*_*h*_, the other positions evolving neutrally. Under this assumption, the expected number of WS and SW substitutions are given by:

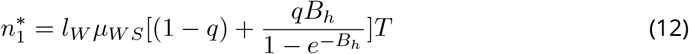

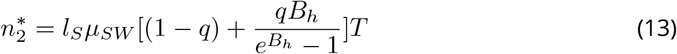

while 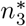. is given by equation 7.

Equations 12 and 13 simplify if *q* is assumed to be much smaller than 1 and *B*_*h*_ much higher than 1; under these assumptions, we have:

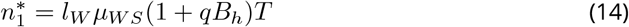

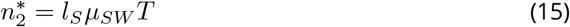

Parameters *q* and *B*_*h*_ only appear as a product in equations 14 and 15, saving one degree of freedom. The simplified equation 15 exhibits the main difference between hotspot and gene-homogeneous models, which concerns SW substitutions. In gene-homogeneous models, the expected number of SW substitutions is decreased in genes experiencing strong gBGC, whereas hotspot models predict nearly no influence of gBGC on the SW substitution rate if *q* is sufficiently small.

The among-gene variation in gBGC strength was here modeled via categories of genes that differed with respect to the prevalence of hotspots, *q*, while sharing the same intensity of gBGC within hotspots, *B*_*h*_. Specifically, we considered a three-category model in which the “coldest” category had no hotspot, i.e., *q*_1_ = 0. The fraction of hotspots in the other two categories, *q*_2_ and *q*_3_, and the relative prevalence of the three categories, *p*_1_, *p*_2_ and *p*_3_, as well as *T* and *B*_*h*_, were free to vary. This model was called M3h (for hotspot); its predictions are given by equations 12, 13 and 7. A simplified hotspot model, M3sh (for simplified hotspot), was also implemented by instead using equations 14, 15 and 7. M3sh is a special case of M3h assuming that the fraction of sites affected by gBGC within a gene is small. We also considered a simplified five-category hotspot model, M5shf, with fixed values for the *q*_*k*_*B*_*h*_ product equal to 0, 0.333, 1, 3.33 and 10, respectively.

The overall likelihood was obtained by multiplying the likelihoods of distinct genes. Parameters were estimated in the maximum likelihood (ML) framework. Likelihood maximization was achieved via home-made C++ programs using the Bio++ library (Guéguen et al., 2013).

The across gene categories average estimated intensity of gBGC was computed as

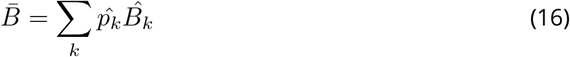

under gene-homogeneous models and

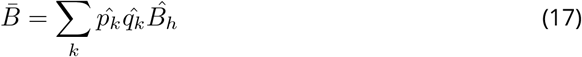

under hotspot models, where the *k* index is for gene categories and the hat denotes ML estimate. The across genes standard deviation of the intensity of gBGC was calculated similarly.

Akaike’s Information Criterion (AIC) was calculated for all models, the number of estimated parameters being 2, 4, 5, 5, 5, 5 and 6, respectively, for M1, M2, M3z, M5f, M3sh, M5shf and M3h. AIC weights (Posada and Buckley, 2004) were used to calculate an across-model estimate of the *B* parameter. The parametrisation of the various models is recapitulated in Table 1.

**Table 1.** Parametrisation of the models used in this analysis. *a* : number of categories of genes; *b*: number of fixed parameters; *c*: list of fixed parameters; *d*: number of optimised parameters; *e*: list of optimised parameters; *f* : parameters shared by all models

| model | cat.<sup>a</sup> | nb fixed<sup>b</sup> | fixed<sup>c</sup> | nb optimised<sup>d</sup> | optimised<sup>e</sup> |
| --- | --- | --- | --- | --- | --- |
| all<sup>f</sup> | | | $\mu_{WS}, \mu_{SW}, \mu_{SS}, \mu_{WW}$ | | $T$ |
| M1 | 1 | 4 | | 2 | $B$ |
| M2 | 2 | 4 | | 4 | $B_1, B_2, p_1$ |
| M3z | 3 | 5 | $B_1$ | 5 | $B_2, B_3, p_1, p_2$ |
| M5f | 5 | 9 | $B_1-B_5$ | 5 | $p_1-p_4$ |
| M3h | 3 | 5 | $q_1$ | 6 | $q_2, q_3, B_h, p_1, p_2$ |
| M3sh | 3 | 5 | $q_1 B_h$ | 5 | $q_2 B_h, q_3 B_h, p_1, p_2$ |
| M5shf | 5 | 9 | $q_1 B_h-q_5 B_h$ | 5 | $p_1-p_4$ |

For each gene, the expected number of WS substitutions in the absence of gBGC was estimated as:

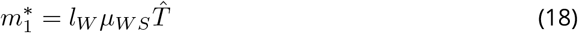

and the excess WS substitutions due to gBGC were estimated as 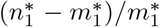. No depletion of SW substitutions is expected under M3sh (compare equations 15 and 18). Finally, each gene was assigned to one of the gBGC categories by selecting the category *k* maximising the following posterior probability:

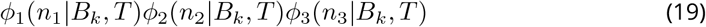

All these calculations were achieved separately for the 40 branches of the data set.

### Additional variables

For every branch, the numbers of non-synonymous substitutions of the WS, SW and SSWW sorts at first and second codon positions were computed, summed across genes, and used to calculate branch-specific *d*_*N*_*/d*_*S*_ ratios. A literature survey was conducted in search for genome-wide estimates of within-species diversity, or heterozygosity, *π*. Such estimates were collected in 18 of the analysed species, as reported in Supplementary Table S2. Data on species longevity and body mass were obtained from the AnAge data base (Magalhães and Costa, 2009) and are also reported in Supplementary Table S2.

## Data accessibility, Supplementary material

All the data sets, programs and scripts used in this study are available from: https://osf.io/fx54q/?view_only=1109ca2f66e74ad99f0d76ac93d40fc5

## Acknowledgements

The author is grateful to Carina Mugal, Fanny Pouyet, David Castellano, an anonymous reviewer, Laurent Duret, Nicolas Lartillot, Sylvain Glémin, Benoît Nabholz and Jonathan Romiguier for very useful comments, and to Adam Eyre-Walker for sharing data on *de novo* mutation in humans. This work was supported by Agence Nationale de la Recherche project ANR-19-CE12-0019 (HotRec). Version 5 of this preprint has been peer-reviewed and recommended by *Peer Community In Genomics* (https://doi.org/10.24072/pci.genomics.100012)

