## Supplementary Figures for "Fine-scale quantification of GC-biased gene conversion intensity in mammals" (from another version of this preprint: version 4, 2021-07-22)

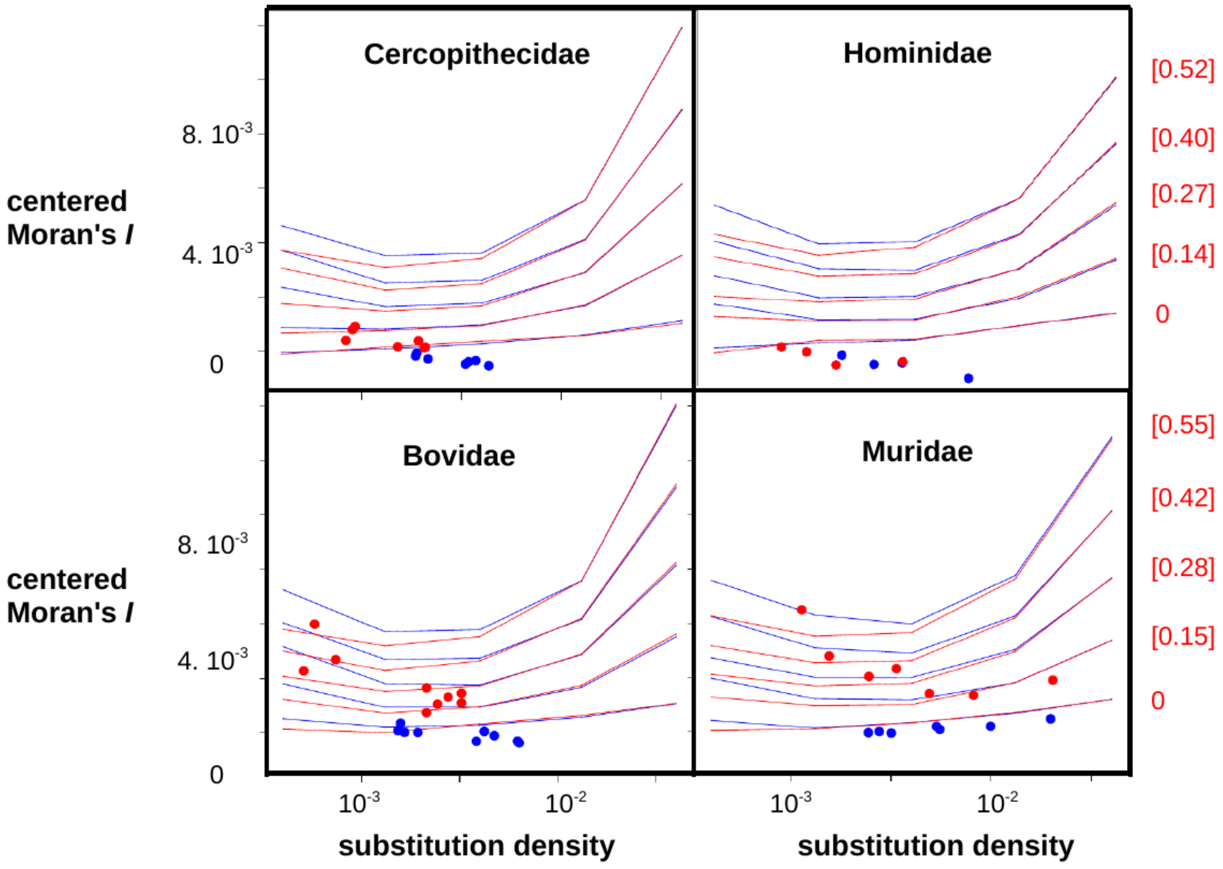


**Supplementary Figure S1. Simulated vs. observed Moran's I**

Blue : SW substitutions ; Red : WS substitutions ; Lines correspond to simulated data, dots to observed data (one dot per lineage; at least 100 genes with >=3 substitutions of each sort required to be included). X-axis : substitutions density (log scale). Y-axis : among genes average centered Moran's I. Substitution density was obtained by dividing the total number of synonymous substitutions by total coding sequence length, then multiplying by 4, such that X-axis is approximately in unit of per synonymous site substitution. There are five lines per substitution sort per panel, which correspond to a prior probability for two successive substitutions to be clustered of 0, 0.1, 0.2, 0.3 and 0.4, respectively. Numbers in red are the realized proportions of clustered substitutions under these conditions, accounting for substitutions that were lost because appearing within introns or flanking regions. These numbers are very similar in Cercopithecidae and Bovidae (not provided due to lack of space).

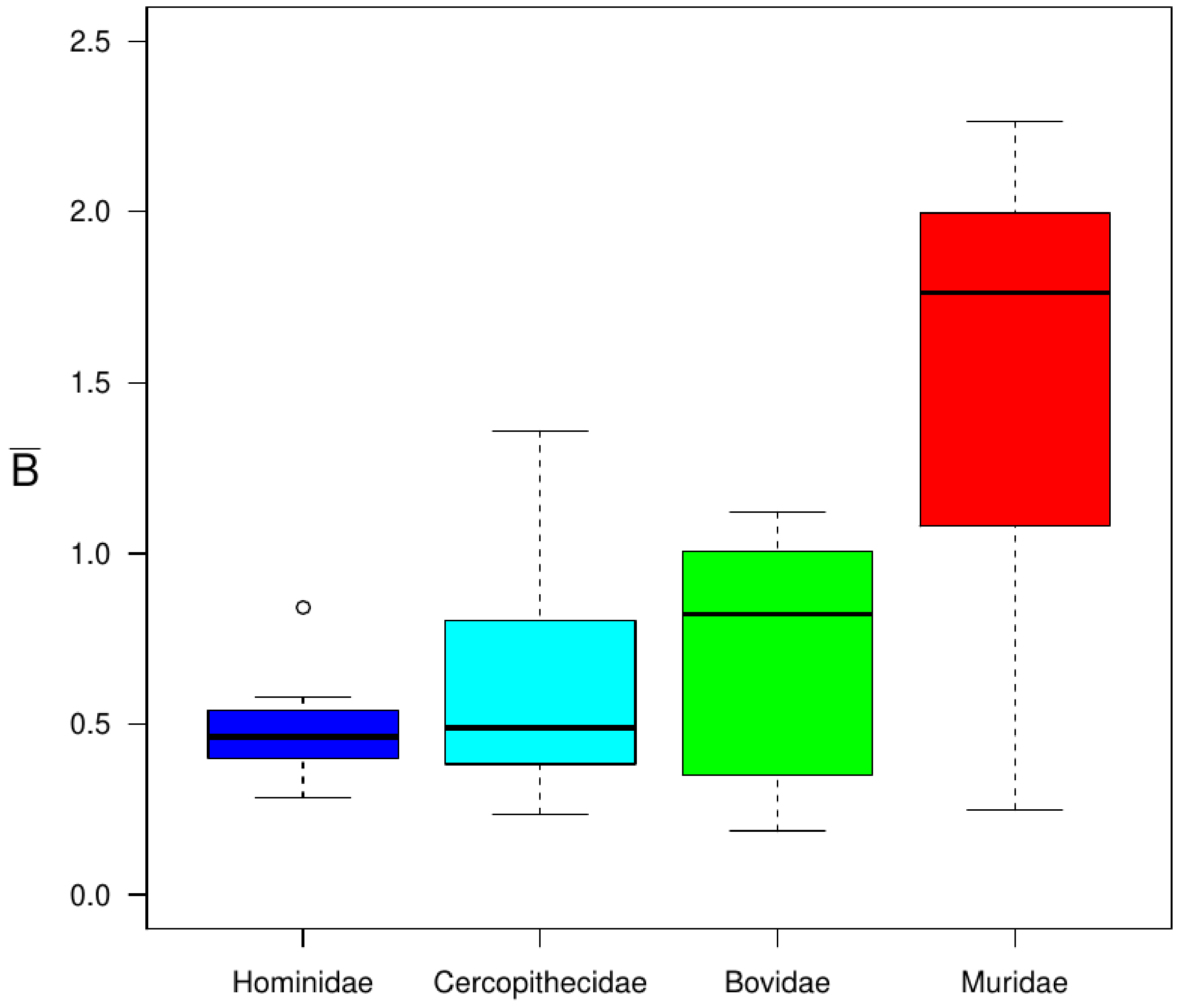


**Supplementary Fig. S2 : Distribution of the average estimated B (Akaike weights).**

Estimates were obtained by taking a weighted average of B estimates under models M1, M2, M3z, M3h and M3sh. For each branch, the weight for model M was equal to exp(-d([M]/2), where d[M] was the difference between the AIC for model M and the minimal AIC across all models. One outlying data point is missing from the figure: the estimated average B was 3.86 in the (*Capra hircus*, *Ovis aries*) ancestral branch (Bovidae).

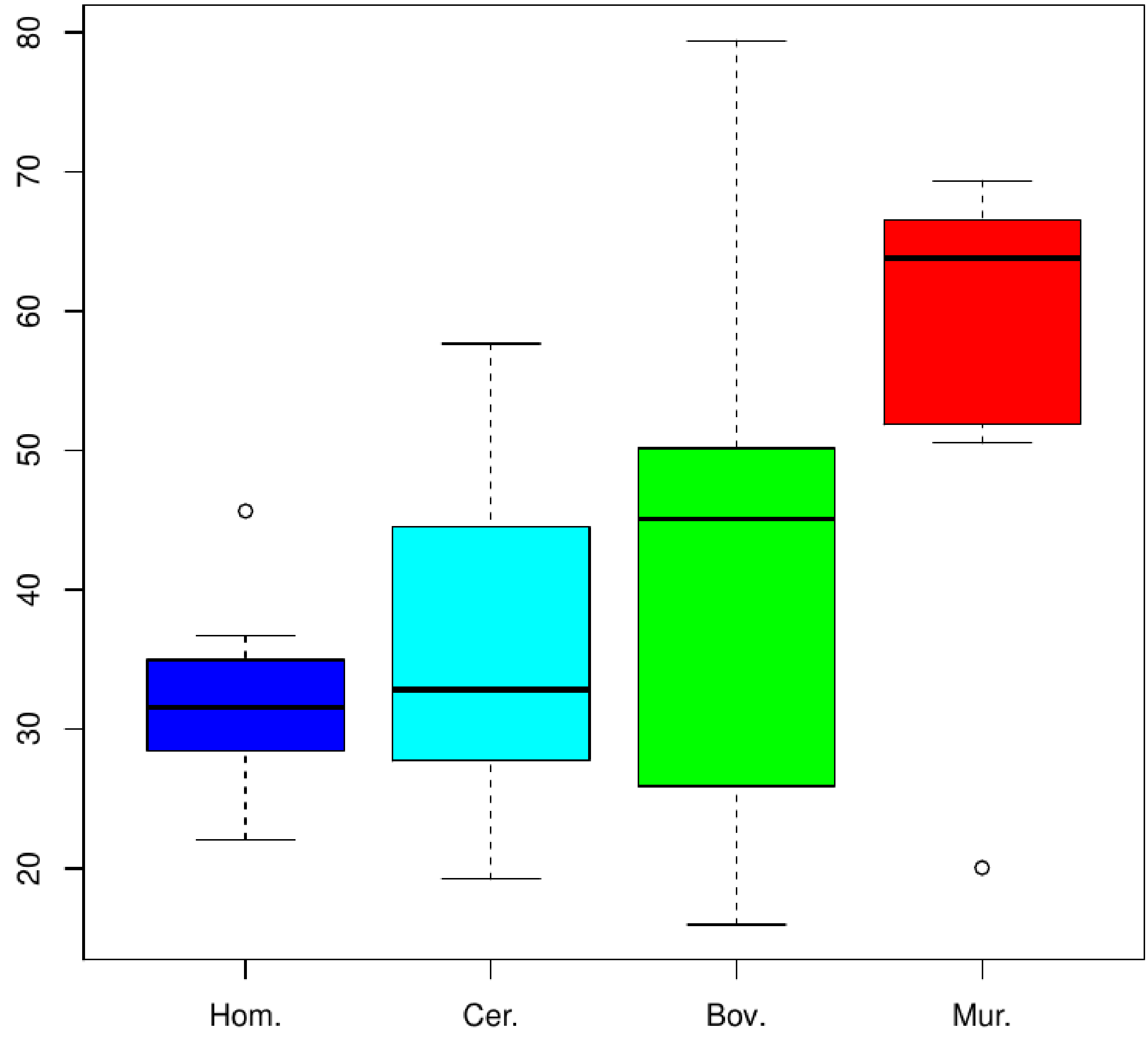


**Supplementary Fig. S3 : Excess WS synonymous substitutions due to gBGC.**

Box plots show the across-lineages distributions of the estimated percentage of WS synonymous substitutions in excess, compared to numbers predicted in the absence of gBGC under model M3sh.

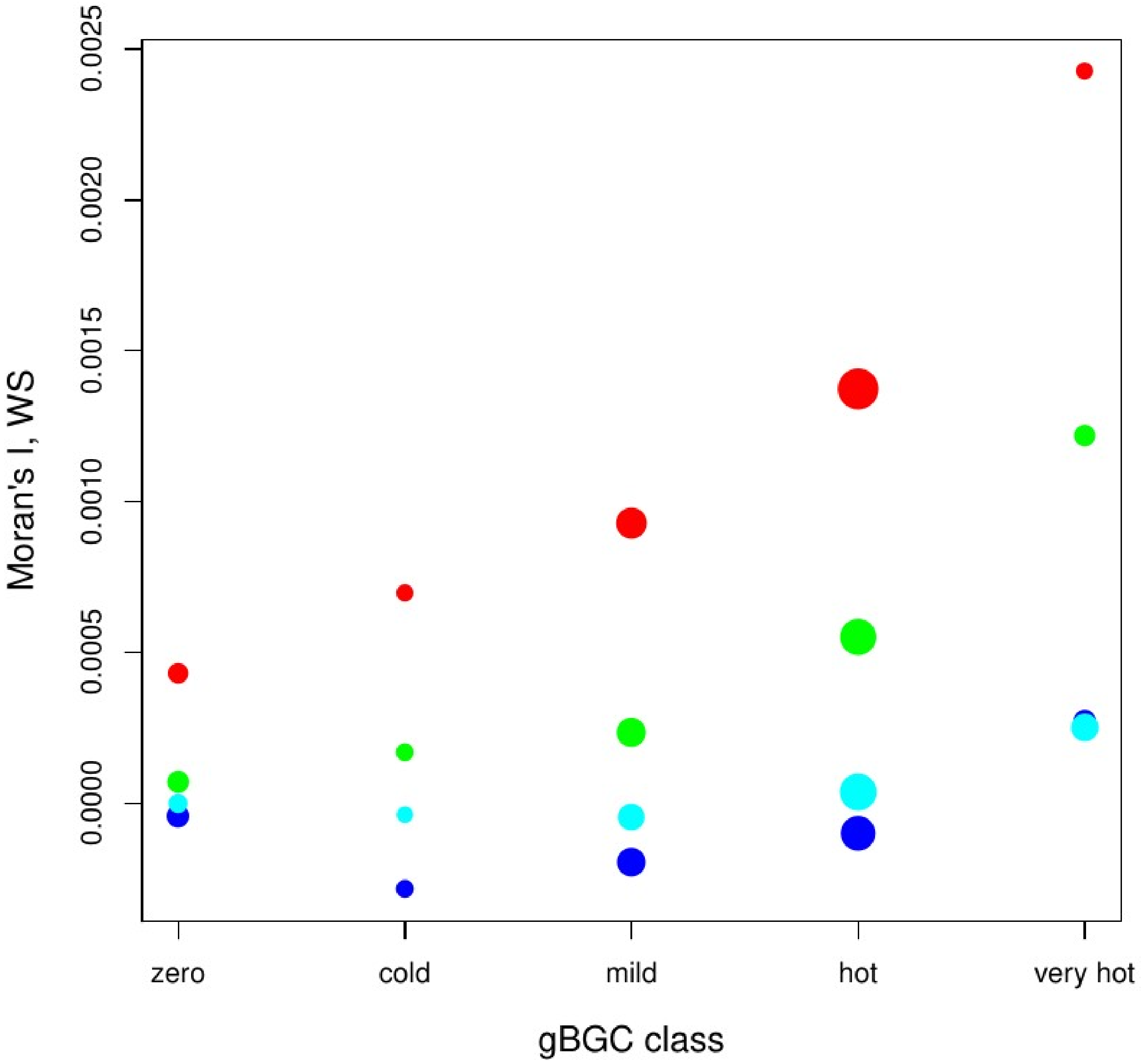


**Supplementary Fig. S4. Variation of Moran's I across gBGC categories of genes and families, model M5shf.**

Legend : see Fig. 5
